# *Candida utilis* yeast as a functional protein source for Atlantic salmon (*Salmo salar* L.): Local intestinal tissue and plasma proteome responses

**DOI:** 10.1101/658781

**Authors:** Felipe Eduardo Reveco-Urzua, Mette Hofossæter, Mallikarjuna Rao Kovi, Liv Torunn Mydland, Ragnhild Ånestad, Randi Sørby, Charles McL. Press, Leidy Lagos, Margareth Øverland

## Abstract

Microbial ingredients such as *Candida utilis* yeast are known to be functional protein sources with immunemodulating effects whereas soybean meal causes soybean meal-induced enteritis in the distal intestine of Atlantic salmon (*Salmo salar* L.). Inflammatory or immunomodulatory stimuli at the local level in the intestine may alter the plasma proteome profile of Atlantic salmon. These deviations can be helpful indicators for fish health and therefore potential tools in diagnosis of fish diseases. The present work aimed to identify local intestinal tissue responses and changes in plasma protein profiles of Atlantic salmon fed inactive dry *Candida utilis* yeast biomass, soybean meal, or combination of soybean meal based diet with various inclusion levels of *Candida utilis*. A fishmeal based diet was used as control diet. Inclusion of *Candida utilis* yeast to a fishmeal based diet did not alter the morphology, immune cell population or gene expression of the distal intestine, but gave a plasma proteome profile different from the fishmeal based control. Lower levels of *Candida utilis* combined with soybean meal modulated immune cell populations in the distal intestine and reduced the severity of soybean meal-induced enteritis, while higher inclusion levels of *Candida utilis* were less effective. The results suggest that *Candida utilis* could induce systemic responses without altering intestinal morphology, and thus could be a high-quality alternative protein source with potential functional properties in diets for Atlantic salmon.

## Introduction

The composition of feeds used in salmon aquaculture has undergone significant changes over the last decades. The rapid growth in the aquaculture industry, but stable production of the major protein resource, fishmeal (FM), has led to increasing use of alternatives. Alternative protein sources are required to contribute to a well-balanced diet and to support optimal fish growth performance, health and disease resistance. Currently, proteins derived from insects [1], terrestrial animal co-products [2, 3] and microbial ingredients [4] are considered to be valuable alternatives. In commercial diets, plant derived proteins have already replaced two third of the proteins of marine origin [5].

Plant ingredients are the most attractive protein sources due to their low cost of production, high protein content and availability [6]. Inclusion of plant ingredients in salmonid feeds can, however, result in reduced growth performance and feed utilization, and health issues due to anti-nutritional factors (ANF) [6, 7]. ANFs in plant-based diets have been associated with detrimental effects on the intestine of several salmonid species [7], and soybean meal-induced enteritis (SBMIE) is a well described condition in Atlantic salmon (*Salmo salar* L.) [8, 9]. Currently, in commercial salmon diets, the refined soy product with reduced level of ANF, soy protein concentrate, is the major protein source of plant origin [5] and has not shown to cause pathological changes in the distal intestine (DI) of salmonids after short-term dietary exposure [9]. However, certain degree of inflammation and ectopic epithelial cells have been observed in the DI of salmonids when fed diets based on FM and soy protein concentrate over a longer period of time [10].

Microbial ingredients have proven to be high quality protein sources with the ability to mitigate the negative effects of plant-derived proteins [11]. Microbial ingredients such as yeast and bacteria contain bioactive compounds with immunemodulating properties that improve the changes caused by SBM [12-14]. Moreover, bacterial meal has been shown to prevent the development of SBMIE in a dose-dependent manner [15]. Mannan oligosaccharides, compounds found in yeast cell walls, have been used as a prebiotic and shown to protect the intestinal mucosa and reduce inflammation and leukocyte infiltration [16]. However, the degree of bioactivity of these compounds depends on the microbial origin as well as the fermentation conditions and downstream processing of the microbial product before incorporation into the salmon diet [11, 17].

Assessing the bioactivity of novel dietary microbial ingredients has traditionally involved the evaluation of local intestinal tissue responses, such as changes in morphology, gene expression or microbiome. Local intestinal responses can induce changes in the whole organism, inducing systemic responses that could contain new biomarkers for health and disease [18]. Innate immune system may respond to local inflammatory or immunomodulatory stimuli in the intestine of fish and in turn elicit changes systemically. Release of cytokines into the circulation stimulates hepatocytes to produce proteins and release them into the circulation to regain homeostasis [19]. Plasma proteomic analysis is a post-genomic tool that allows investigation of complex biological systems involved in physiology and pathology. Plasma proteome profiles in response to certain inflammatory or immunomodulatory stimuli can be useful diagnostic tools for fish diseases and indicators of fish health.

In this study, an inactive dry yeast strain *Candida utilis* (*C. utilis*) was used as an alternative protein source with functional properties in FM and SBM based diets. SBM was used as a dietary challenge to induce SBMIE. Increasing levels of *C. utilis* were included in the diets to evaluate the immunemodulating properties of the yeast, in particular the ability to counteract SBMIE. We combine histomorphological evaluation, immunohistochemistry, morphometry and gene expression of the DI with plasma proteome analysis. By combining these methods, we aim to identify local intestinal tissue responses and changes in plasma protein profiles in Atlantic salmon resulting from dietary treatments. Our results show that inclusion of *C. utilis* as an alternative protein source could induce a systemic response in plasma proteins without altering the local morphology and immune cell population in the DI of Atlantic salmon.

## Materials and methods

### Experimental ingredient and diet preparation

The inactive dry *C. utilis* yeast corresponded to a commercial product called Lakes States® Type B produced by LALLEMAND SAS (Blagnac, France). Supplementary Table 1 (S1 Table) shows the proximate composition of the test inactive dry yeast. The crude protein content was determined by multiplying nitrogen content by a conversion factor of 6.25. All diets used in this study were formulated to meet or exceeded the nutrient requirements of Atlantic salmon [20] (Table 1), produced by extrusion technology at the Center for Feed Technology (FôrTek) at the Norwegian University of Life Sciences (Aas, Norway), and stored at −20°C prior to feeding. The extruded pellets were dried to ∼ 6 % moisture content before vacuum coating with fish oil. The diets consisted of a FM-based control diet (FM group) and the following six experimental diets; a diet containing 200 g/kg *C. utilis* (FM200CU group), and five diets containing 200 g/kg SBM together with 0 (SBM group), 25, 50, 100 or 200 g/kg *C. utilis* (SBM25CU, SBM50CU, SBM100CU and SBM200CU groups, respectively).

**Table 1.**
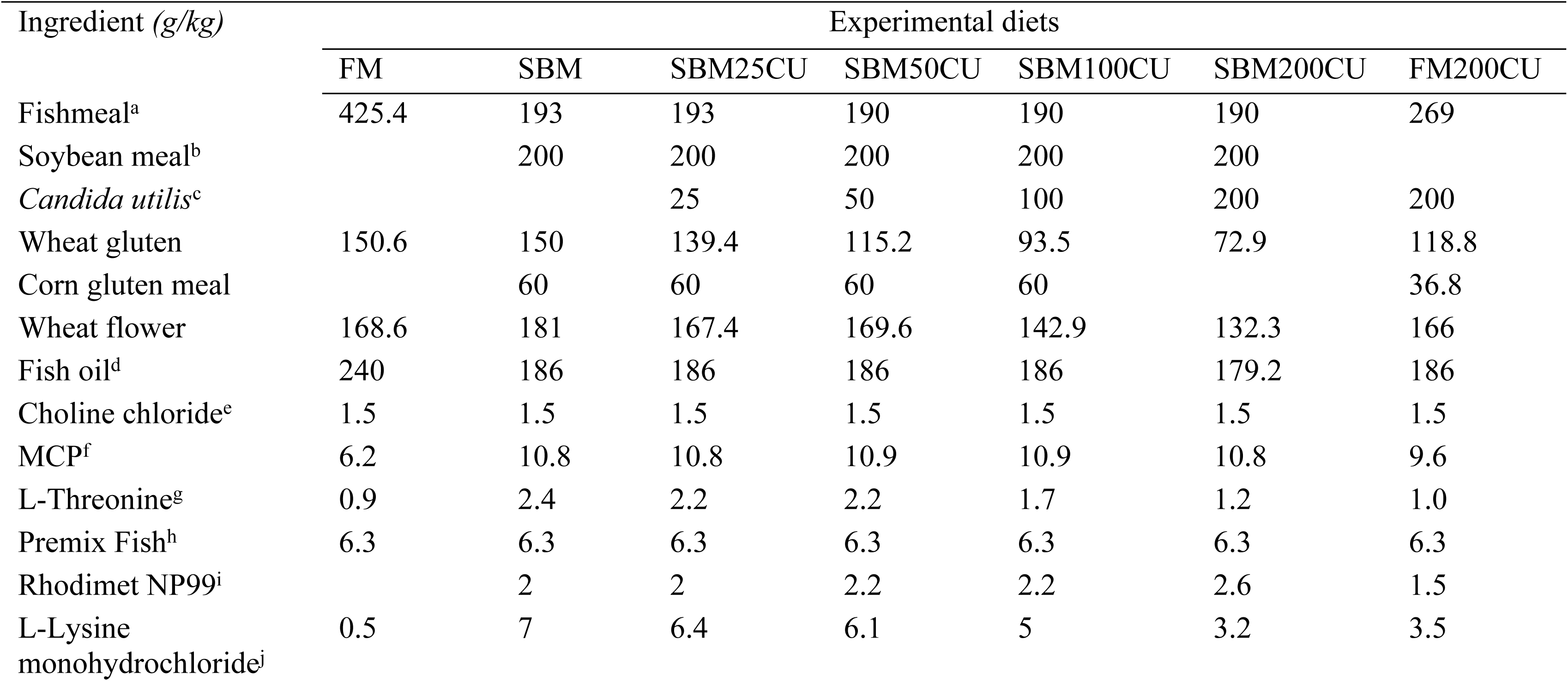
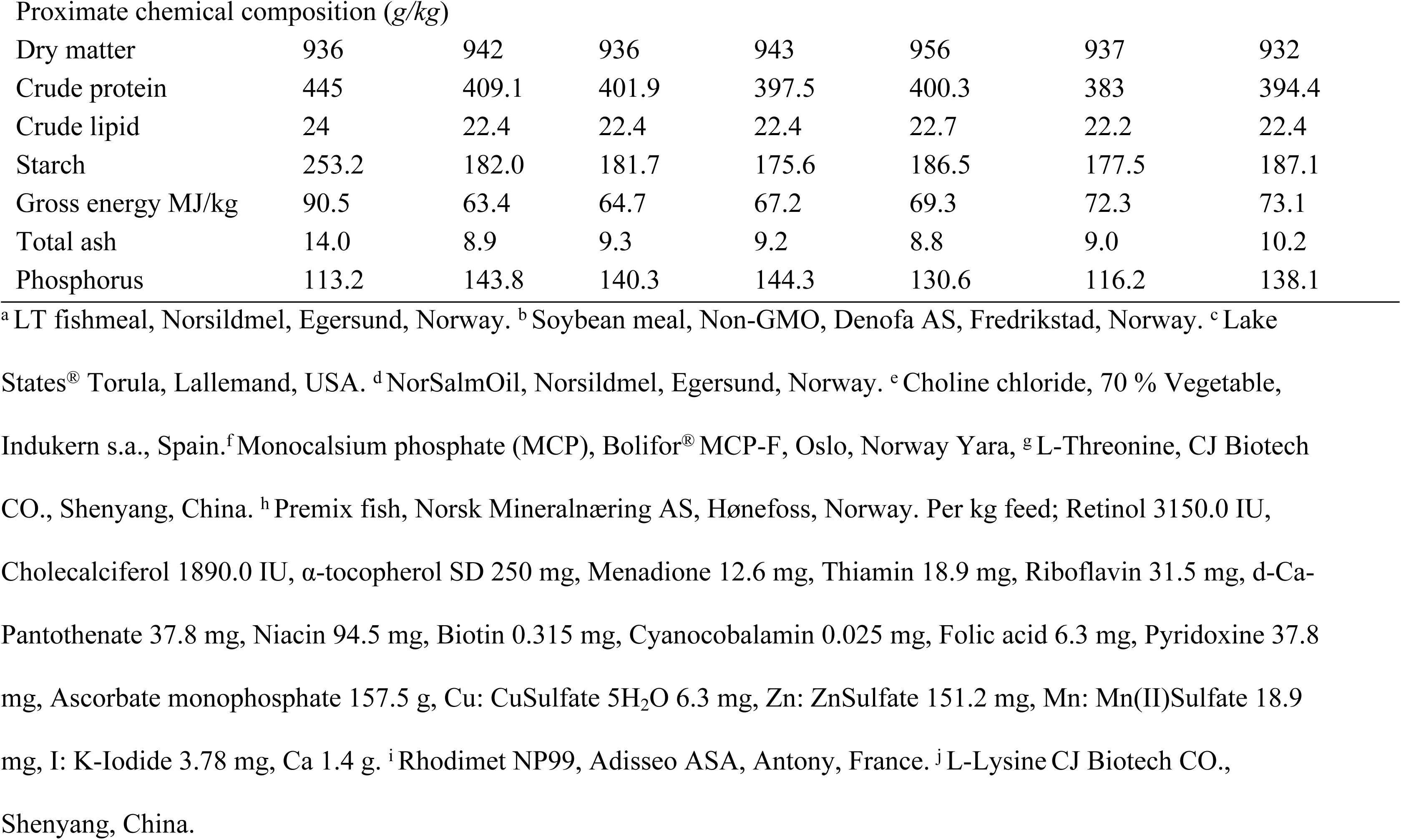
Ingredient and proximate chemical composition (g/kg) of control (FM) and experimental diets. FM = Fishmeal; SBM = soybean meal; SBM25CU = soybean meal + 25 g/kg *C. utilis*; SBM50CU = soybean meal + 50 g/kg *C. utilis;* SBM100CU = soybean meal + 100 g/kg *C. utilis*; SBM200CU = soybean meal + 200 g/kg *C. utilis*; FM200CU = fishmeal + 200 g/kg *C. utilis*.

| Ingredient (g/kg) | Experimental diets |  |  |  |  |  |  |
| --- | --- | --- | --- | --- | --- | --- | --- |
|  | FM | SBM | SBM25CU | SBM50CU | SBM100CU | SBM200CU | FM200CU |
| Fishmeal<sup>a</sup> | 425.4 | 193 | 193 | 190 | 190 | 190 | 269 |
| Soybean meal<sup>b</sup> |  | 200 | 200 | 200 | 200 | 200 |  |
| *Candida utilis*<sup>c</sup> |  |  | 25 | 50 | 100 | 200 | 200 |
| Wheat gluten | 150.6 | 150 | 139.4 | 115.2 | 93.5 | 72.9 | 118.8 |
| Corn gluten meal |  | 60 | 60 | 60 | 60 |  | 36.8 |
| Wheat flower | 168.6 | 181 | 167.4 | 169.6 | 142.9 | 132.3 | 166 |
| Fish oil<sup>d</sup> | 240 | 186 | 186 | 186 | 186 | 179.2 | 186 |
| Choline chloride<sup>e</sup> | 1.5 | 1.5 | 1.5 | 1.5 | 1.5 | 1.5 | 1.5 |
| MCP<sup>f</sup> | 6.2 | 10.8 | 10.8 | 10.9 | 10.9 | 10.8 | 9.6 |
| L-Threonine<sup>g</sup> | 0.9 | 2.4 | 2.2 | 2.2 | 1.7 | 1.2 | 1.0 |
| Premix Fish<sup>h</sup> | 6.3 | 6.3 | 6.3 | 6.3 | 6.3 | 6.3 | 6.3 |
| Rhodimet NP99<sup>i</sup> |  | 2 | 2 | 2.2 | 2.2 | 2.6 | 1.5 |
| L-Lysine monohydrochloride<sup>j</sup> | 0.5 | 7 | 6.4 | 6.1 | 5 | 3.2 | 3.5 |
| Dry matter | 936 | 942 | 936 | 943 | 956 | 937 | 932 |
| Crude protein | 445 | 409.1 | 401.9 | 397.5 | 400.3 | 383 | 394.4 |
| Crude lipid | 24 | 22.4 | 22.4 | 22.4 | 22.7 | 22.2 | 22.4 |
| Starch | 253.2 | 182.0 | 181.7 | 175.6 | 186.5 | 177.5 | 187.1 |
| Gross energy MJ/kg | 90.5 | 63.4 | 64.7 | 67.2 | 69.3 | 72.3 | 73.1 |
| Total ash | 14.0 | 8.9 | 9.3 | 9.2 | 8.8 | 9.0 | 10.2 |
| Phosphorus | 113.2 | 143.8 | 140.3 | 144.3 | 130.6 | 116.2 | 138.1 |

<sup>a</sup> LT fishmeal, Norsildmel, Egersund, Norway. <sup>b</sup> Soybean meal, Non-GMO, Denofa AS, Fredrikstad, Norway. <sup>c</sup> Lake
States<sup>®</sup> Torula, Lallemand, USA. <sup>d</sup> NorSalmOil, Norsildmel, Egersund, Norway. <sup>e</sup> Choline chloride, 70 % Vegetable,
Indukern s.a., Spain. <sup>f</sup> Monocalcium phosphate (MCP), Bolifor<sup>®</sup> MCP-F, Oslo, Norway Yara, <sup>g</sup> L-Threonine, CJ Biotech
CO., Shenyang, China. <sup>h</sup> Premix fish, Norsk Mineralnæring AS, Hønefoss, Norway. Per kg feed; Retinol 3150.0 IU,
Cholecalciferol 1890.0 IU, $\alpha$ -tocopherol SD 250 mg, Menadione 12.6 mg, Thiamin 18.9 mg, Riboflavin 31.5 mg, d-Ca-
Pantothenate 37.8 mg, Niacin 94.5 mg, Biotin 0.315 mg, Cyanocobalamin 0.025 mg, Folic acid 6.3 mg, Pyridoxine 37.8
mg, Ascorbate monophosphate 157.5 g, Cu: CuSulfate 5H<sub>2</sub>O 6.3 mg, Zn: ZnSulfate 151.2 mg, Mn: Mn(II)Sulfate 18.9
mg, I: K-Iodide 3.78 mg, Ca 1.4 g. <sup>i</sup> Rhodimet NP99, Adisseo ASA, Antony, France. <sup>j</sup> L-Lysine CJ Biotech CO.,
Shenyang, China.

### Fish husbandry and feeding trial

Vaccinated salmon were acquired from Sørsmolt AS (Sannidal, Norway) and maintained according to the guidelines established by the Norwegian Animal Research Authority at the Research Station Solbergstrand of Norwegian Institute of Water Research (Drøbak, Norway). Fish were acclimated to seawater, housed in 300 L tanks supplied with ultraviolet light treated seawater (8 °C; 34.5 g/L NaCl) in a 7–8 L per min flow-through system, and fed with a commercial marine-based compound feed not containing soybean-derived products (3-mm pellet; Polarfeed AS, Europharma, Leknes, Norway) under continuous light during a 4-month period prior to conducting the feeding trial. The water temperature, dissolved oxygen and pH level were measured and recorded daily. At the beginning of the experiment, 360 fish (average initial body weight of 526 g) were randomly assigned to 18 tanks (20 fish/tank) and acclimated to the FM based control diet for two weeks prior to feeding experimental diets. Feeding was approximately 20% in excess twice daily using automatic belt feeders based on a daily estimate of fish biomass and uneaten feed per tank, which was collected from the tank outlet after each feeding period. Following the acclimation period, each experimental diet was randomly allocated to the fish tanks (two tanks/diet) and fed for 30 days as described above. During the experiment, fish groups were batch weighed at 0, 7, 30 and 37 days and individual weight and length of sampled fish were recorded. After 30 days, the feeding strategy were changed and fed for 7 days as illustrated in Fig 1.

**Fig 1.**
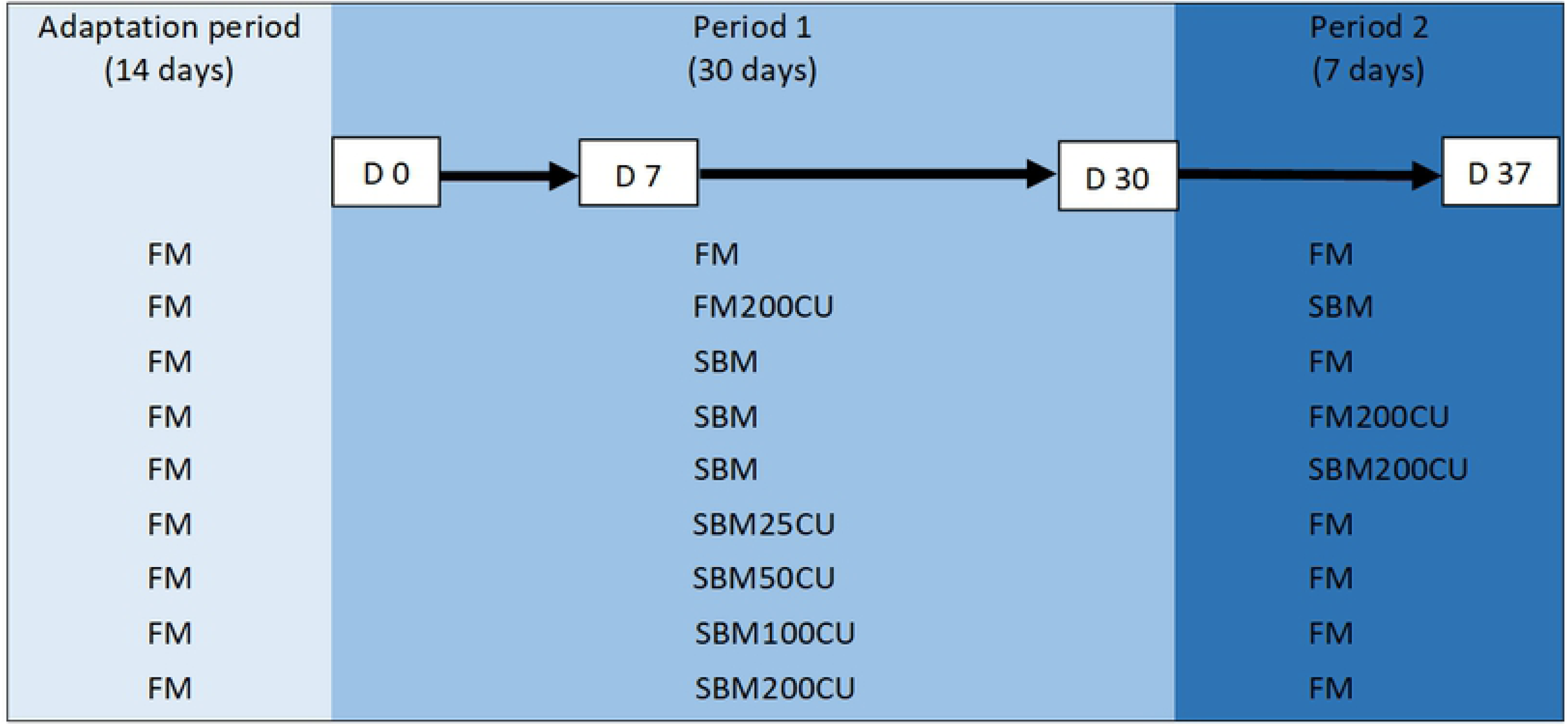
Experimental design. The adaptation period of 14 days was followed by experimental period 1 that lasted for 30 days. In experimental period 2, there was a shift in diets and this period lasted for 7 days. Sampling points are day 0, 7, 30 and 37. FM = Fishmeal; SBM = soybean meal; SBM25CU = soybean meal + 25 g/kg *C. utilis*; SBM50CU = soybean meal + 50 g/kg *C. utilis;* SBM100CU = soybean meal + 100 g/kg *C. utilis*; SBM200CU = soybean meal + 200 g/kg *C. utilis*; FM200CU = fishmeal + 200 g/kg *C. utilis*.

### Sample collection

At each sampling point, 8 fish per diet (4 fish per tank) were randomly selected and anaesthetized by immersion in 60 mg/l of tricaine methanesulfonate (MS-222, Sigma-Aldrich, MO, USA) and subsequently euthanized by a sharp blow to the head prior to dissection at the following sampling points: 0, 7, 30 and 37 days after starting feeding experimental diets. Before dissecting fish, blood samples were taken from the *vena caudalis* (tail vein) using heparinized syringe and centrifuged (1300-2000 x g for 10 min) to isolate blood plasma, which was aliquoted and stored at –80°C, until proteomic analysis was performed. DI tissue was sampled and kept in RNAlater® (Merck KGaA, Darmstadt, Germany) at 4°C for 24 h, and then at –80°C, until RNA extraction. DI tissue samples were also collected and preserved for subsequent histology, morphometry and immunohistochemistry.

### Histology, immunohistochemistry and morphometric measurements Histology

Approximately 1 cm segment of the DI was open longitudinally and the intestinal content were carefully removed. The tissue was fixed in 10% formalin for 48 h at room temperature and further processed according to routine histological procedures. Briefly, tissue was embedded in paraffin with an orientation to ensure longitudinal sectioning. Sections (2 µm) of paraffin-embedded DI tissue were mounted on glass slides (Menzel Gläser, Thermo Fisher scientific, Braunschweig, Germany) and processed for staining with hematoxylin and eosin. A blinded, semi-quantitative histological scoring of the DI was performed using the criteria described in detail by Baeverfjord and Krogdahl [8]. Briefly, the criteria were: 1) shortening of both the simple and complex intestinal mucosal folds, 2) appearance of the enterocytes including supranuclear vacuolization, cellular heights and presence of intraepithelial lymphocytes (IELs), 3) widening of the central lamina propria of the simple and complex folds by connective tissue and 4) infiltration of leucocytes in the lamina propria. Each criterion was given a score ranging from 0 to 2, and half scores were included (i.e. 0, 0.5, 1, 1.5, 2) [13]. Score 0 indicated normal morphology and score 2 represented marked changes. Score 0.5 was regarded as changes within normal range.

### Immunohistochemistry

Fish subjected for immunohistochemical analysis were from sampling at day 30, and the following diet groups were included: FM, FM200CU, SBM, SBM25CU and SBM200CU. CD3 and CD8 positive T lymphocytes were identified in DI tissue sections by immunohistochemistry using a monoclonal anti-CD3ε antibody (dilution 1:600) [21] and a monoclonal anti-CD8α antibody (kindly supplied by Karsten Skjødt, dilution 1:50) [22], respectively. This analysis was performed as follows: formalin-fixed, paraffin embedded DI sections (4 µm) were mounted on poly-L-lysine-coated glass slides (Superfrost Plus, Thermo Fisher scientific, Braunschweig, Germany) and left to dry at 37°C. The slides were incubated at 58°C for 30 min before deparaffinised in xylene and rehydrated in graded alcohols to distilled water. Antigen retrieval was done by using hydrated autoclaving at 121°C for 15 min in 0.01 M citrate buffer, pH6. Endogenous peroxidase was inhibited with 0.05 % phenylhydrazine (0.05%; Sigma-Aldrich, MO, USA) in phosphate buffered saline, preheated to 37°C, for 40 min. The sections were stored in PBS overnight at 4°C and then incubated with normal goat serum (dilution 1:50; Sigma-Aldrich) in 5 % bovine serum albumin /0.05 M tris-buffered saline to avoid non-specific binding for 20 min. The blocking solution was tapped off without washing, and the sections were incubated with primary antibody diluted in 1% bovine serum albumin/Tris-buffered saline for 1 h. Control sections were incubated with only 1% bovine serum albumin. The sections were incubated in the kit polymer-HRP anti-mouse (Dako En Vision+ System-HRP, Dako, Glostrup, Denmark), as secondary antibody, for 30 min. Peroxidase activity was detected with 3,3’-diaminobenzidine following the kit procedure. The sections were counterstained with haematoxylin for 30 s followed by washing in distilled water before mounting with Aquatex (Novoglas Labortechnik Langenbrinck, Bern, Germany). Unless otherwise stated, the sections were washed three times with phosphate buffered saline for 5 min between each step. All incubations took place in a humid chamber at room temperature.

### Morphometric measurements and calculation of immune cell density

Morphometric measurements and calculation of the density of immune cells were performed from immunohistochemically labelled sections mentioned above. ImageJ software, version v1.51r [23], was used to perform the measurements and calculations. Images were captured with a Zeiss Axiocam 506 color camera connected to a light microscope (Zeiss Axio Imager M2, Carl Zeiss, Germany) at a 10 X magnification. The measurement scale was set to 2.26 pixels/µm in ImageJ and the measurements were converted from µm to mm. Counting of immunohistochemically labelled cells was performed using the multi point tool. The freehand selection and segmented line selection were used to measure fold area and height, respectively. The fold height was measured from stratum compactum to the tip of the epithelium lining the fold (S1 Fig.). The fold area was measured from stratum compactum, including the middle of the fold base on each side, and the whole simple fold (S1 Fig.). The immunohistochemically labelled cells were counted within the measured area of the simple fold. The density of CD3ε- or CD8α-labelled cells was calculated as follow: Cell density = (no. of labelled cells)/area. Simple folds were subjected to all the measurements mentioned above and the folds were selected as the first appropriate simple fold located to the left of a complex fold. An appropriate fold was defined as a fold that appeared long, not bent and had an intact epithelium that was attached to the basement membrane all the way to the tip of the fold. Between 2 - 6 measurements were collected from each individual with a total of at least 30 measurements from each group for each measurement. A mean for each individual was calculated based on the measurements.

### RNA isolation, cDNA synthesis, quantitative PCR

DI from FM, FM200CU, SBM and SBM200CU diet groups (8 fish/diet) at day 30 were subject to gene expression analysis. Total RNA was extracted and purified using RNeasy® 96 kit (Qiagen, Valencia, USA) and QIAcube® HT system (Qiagen), according to the manufacturer’s protocol. After the first washing step, on-column DNase treatment was performed using PureLink TM DNase kit (Thermo Fisher Scientific, Waltham, Massachusetts). RNA concentration and quality were measured using NanoDrop TM 8000 spectrophotometer (Thermo Fisher Scientific). Purified total RNA was stored at −80°C until further analysis.

Prior to cDNA synthesis, all samples were normalized to 400 ng/µL. cDNA synthesis was performed using AffinityScript QPCR cDNA Synthesis kit following the manufacturer’s guidelines (Agilent Technologies, Santa Clara, CA, USA). The total reaction volume was 10 µL using 5 µL of Mastermix, 1.5 µL random hexamer primers, 0.5 µL AffinityScript RT/ RNase Block enzyme mixture and 3 µL DNase treated RNA. The resulting cDNA was stored at −80 °C before use.

All quantitative PCR (qPCR) reactions were performed in duplicates and conducted in 96 well plates on LightCycler® 480 system (Roche Diagnostics, Mannheim, Germany). Each reaction consisted of a total amount of 12 µL divided into 6 µL LightCycler 480 SYBR Green I Master (Roche Diagnostics), 2 µL primers, and 4 µL cDNA. The qPCR conditions were 95°C for 5 min before a total of 45 cycles of 95°C for 5 seconds, 60°C for 15 seconds, and 72°C for 15 seconds. To confirm amplification specificity, each PCR product was subject to melting curve analysis (95°C 5 s; 65°C 60 s; 97°C continuously). Primers tested are listed in Supplementary Table 2 (S2 Table). Glyceraldehyde-3-phosphate dehydrogenase (GDPH) and hypoxanthine phosphoribosyltransferase I (HPRTI) were chosen as reference for normalization since these genes did not show significant differential expression between the diets and have previously been described as suitable reference genes in the DI of salmon [24]. The crossing point (Cp) values were determined using the maximum second derivative method on the basis of the LightCycler® 480 software release 1.5.1.62 (Roche Diagnostics). The geometric mean of the CP-values for GDPH and HPRTI was used an index. The qPCR relative expression of mRNA was calculated using the ΔΔCt method [25].

### In-solution digestion and protein sequence analysis by LC-MS/MS

Proteomic analysis was performed, according to methods described by Lagos *et al.* 2017 [26], using four biological replicates per treatment of plasma taken at the end of the first feeding period (30 days) (FM, SBM, SBM200CU and FM200CU). In brief, frozen plasma (−80° C) was thawed and diluted to 40 µg of total protein in PBS, and the pH was adjusted to 8 by adding ammonium bicarbonate (Sigma-Aldrich, Darmstadt, Germany). The samples were then subjected to overnight incubation at 37°C. The tryptic peptides were dissolved in 10 µL 0.1% formic acid/2% acetonitrile and 5 µL analyzed using an Ultimate 3000 RSLCnano-UHPLC system connected to a Q Exactive mass spectrometer (Thermo Fisher Scientific, Bremen, Germany) equipped with a nano electrospray ion source. The proteomic analysis was performed by the Proteomic core facility of University of Oslo. The mass spectrometry proteomics data have been deposited to the ProteomeXchange Consortium via the PRIDE [27] partner repository with the dataset identifier PXD012051.

## Data analysis

### Histology, morphometric measurements, T-cell density and gene expression

Non-parametric data from the histological evaluation were analyzed by Kruskal-Wallis followed by post hoc Dunn’s test with comparison of mean rank using GraphPad Prism, version 7.00 (GraphPad Software Inc., San Diego, CA, USA). Morphometric analyses and T-cell density were analyzed by one-way ANOVA followed by Tukey-Kramer HSD by using JMP statistical software (JMP®Pro 13.0.0, SAS Institute Inc, NC, USA). Morphometric analyses and T-cell density analyses were performed at the individual level using the mean of measurements of between 2-6 simple folds per fish. Results of qPCR (means ± standard deviations) were analyzed using One-way ANOVA with Dunnett’s multiple comparison test (*a* < 0.0001) on GraphPad Prism 8.0. 1.

### Proteomic data analysis

The resulting proteomic data, MS raw files, were analyzed using MaxQuant and identifications were filtered to achieve a protein false discovery rate (FDR) of 1%. Raw data files were processed using Scaffold4 (Proteome Software, Portland, OR, USA), and a non-redundant output file was generated for protein identifications with log (e) values less than −1. Peptide identification was determined using a 0.8 Da fragment ion tolerance. MS/MS spectra were searched against the salmon proteome, and reverse database searches were used in estimation of false discovery rates. The analysis was restricted to proteins reproducibly identified in at least two of the four replicates per diet, making the minimum number of peptides used to identify each protein an average value of 2. Row-wise normalization was applied to provide Gaussian-like distributions [28] for adjusting the differences among protein data. Protein raw data transferred to log normalization and then performed on autoscaled data (mean-centered and divided by the standard deviation of each variable) [29]. A diagnostic plot was utilized to represent normalization procedures for normal distribution assessments [30]. Spearman’s correlation analysis, multivariate statistical analysis and data modeling were performed in R, using R package MetaboAnalystR [28]. In hierarchical cluster analysis, each individual begins as a separate cluster and the algorithm proceeds to combine them until all individuals belong to one cluster. The clustering results are also presented in the form of a heatmap, with levels of protein expression across the dietary groups. Hierarchical clustering was performed with the hclust function in R package stat. UniprotKB database was used for functional annotation of the proteins. A gene ontology (GO) analysis on the proteomic data was performed to provide insights into the complex structure and biological processes of the plasma proteome across the diets.

A series of principal component analysis (PCA) was performed using the prcomp package in R. PCA was used as an unsupervised method to find the directions of maximum covariance among FM, SBM, SBM200CU and FM200CU. In addition, a series of Partial least squares discriminant analysis (PLS-DA) regression was performed using the plsr function provided by R pls package. The classification and cross-validation were performed using the corresponding wrapper function by the caret package in R. Briefly, to define the optimal number of PCs (principal components), “7-fold cross-validation” (CV) was applied[31]. Using CV, the predictive power of the model was verified. Two parameters were calculated for evaluating the models: R2 (goodness of fit) and Q2 (goodness of prediction). A model with Q2>0.5 is considered good, Q2>0.9 excellent [32]. As cross-validation only assesses the predictive power without a statistical validation, the performance of PLS-DA models was also validated by a permutation test (200 times).

In order to help interpretation of results from PLS-DA, we considered the variable importance in the projection (VIP) scores. This allowed visualization of protein influence (including prediction performance) on the model and identified the best descriptors of the differences among the four conditions. The VIP score is a weighted sum of squares of the PLS loading weights taking into account the amount of explained Y-variation in each dimension [33]. VIP scores “greater than 2” rule is generally used as a criterion to identify the most significant variables [33].

## Results

### Growth and general health

All groups of fish accepted their allocated diets and no significant differences were found in feed intake among dietary treatment. The average initial weight was 526 g and average final weight was 667 g at day 37, indicating that all fish gained weight during the experimental period. The general health of the fish in this experiment was good, but two fish died during the experimental period, one from SBM50CU group and one from SBM100CU group, for unknown reasons.

### Histology

All fish sampled at day 0 showed normal DI morphology. Briefly, simple folds were long and slender with a thin lamina propria, whereas the complex folds were tall with a narrow lamina propria and a partial central core of smooth muscle. Intestinal epithelial cells were tall with the nucleus located basally, large vacuoles located apically and many IELs. Goblet cells were scattered among the epithelial cells towards the apex, and there was a higher presence of goblet cells at the apex of the complex folds. The lamina propria adjacent to the stratum compactum was thin and numerous eosinophilic granule cells were present (data not shown).

At day 7 (Fig 2A), FM and FM200CU groups showed normal DI morphology as described above. In general, all fish groups fed diets containing SBM, including SBM diets with *C. utilis* inclusion, displayed changes in the DI morphology consistent with SBMIE (described in detailed below). Nevertheless, in SBM25CU and SBM50CU, there was variation within the groups ranging from individuals showing no changes to other individuals with moderate changes in DI morphology.

**Fig 2.**
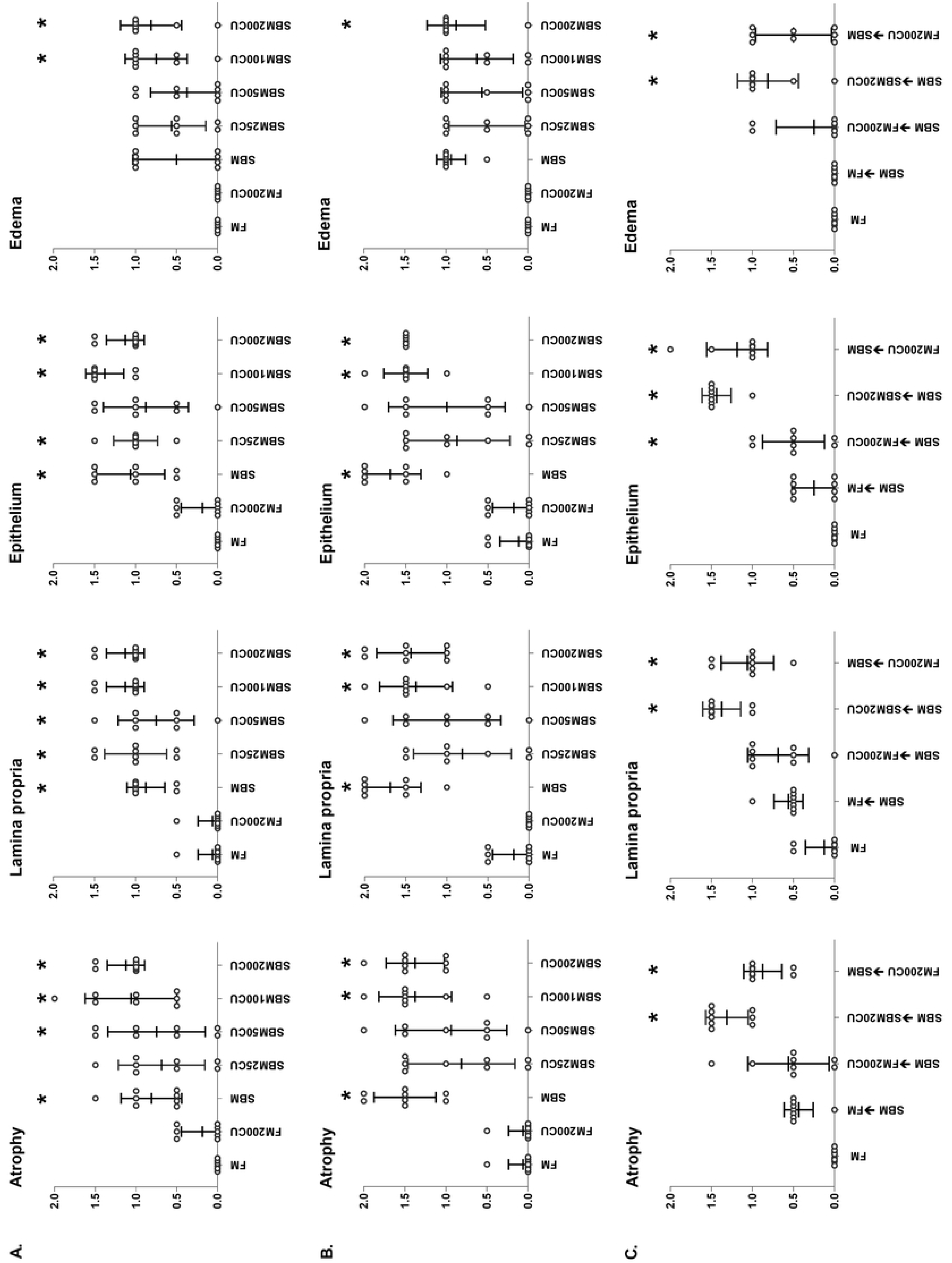
Histological evaluation. Histological evaluation of the distal intestine of Atlantic salmon based on atrophy, lamina propria, epithelium and edema at 7 (A), 30 (B) and 37 (C) days. Changes are scored from 0 to 2 where 0 indicates no changes and 2 indicates severe changes. Data are expressed as mean and standard deviation, n = 8 for all groups. Significant difference from the control fish fed FM based diet is denoted by asterisk (*) (p<0.05; Dunn’s test). FM = fishmeal; FM200CU = fishmeal combined with 200 g/kg *C. utilis* (CU); SBM = soybean meal; SBM25CU = soybean meal with 25 g/kg CU; SBM50CU = soybean meal with 50 g/kg CU; SBM100CU = soybean meal with 100 g/kg CU; SBM200CU = soybean meal with 200 g/kg CU.

At day 30 (Fig 2B), no morphological changes were seen in the DI of FM and FM200CU groups. However, a moderate SBMIE was present in the DI of SBM, SBM100CU and SBM200CU groups. Both simple and complex folds were shorter with a widening of the lamina propria within the folds and adjacent to the stratum compactum. Fusion of the simple folds was frequently observed. There was an increased presence of connective tissue in the lamina propria and the increased infiltration of leucocytes consisted mainly of eosinophilic granule cells and to a lesser extent of lymphocytes, some macrophages and neutrophils were present. The intestinal epithelial cells shown a reduction in height with nucleus displaced in a more apical position and small supranuclear vacuoles. There was an increased presence of IELs. In SBM25CU and SBM50CU groups, there were still a variation within the groups as seen at day 7.

At day 37 (Fig 2C), fish previously fed SBM, either alone or in combination with *C. utilis*, had normal DI morphology after being fed FM for 7 days. Similarly, most of the fish that had diets changed from SBM to FM200CU had DI morphology regarded as normal, but there were some fish in these groups that had a mild enteritis. Changing diet from FM200CU to SBM induced in general a mild SBMIE, whereas a shift from SBM to SBM200CU maintained a moderate SBMIE. The FM control group was normal, as described above.

No tank effects were observed at any of the sampling points. In Fig 2A and 2B, only fish from the tanks that were subjected to immunohistochemical analysis at day 30 are presented, i.e., two SBM-groups are omitted from the figure. Fish with diet shift from SBM combined with *C. utilis* to FM at day 37 had normal DI morphology but are excluded from Fig 2C as this group did not differ from the SBM group.

### Immunohistochemistry, morphometry, density of immune cells

At day 30, CD3ε positive cells showed an abundant presence at the base of the epithelium and extending along the entire length of simple folds of FM and FM200CU (Fig 3A and 3B) groups. Only a few CD3ε positive cells were observed in the lamina propria adjacent to the stratum compactum, and were rarely present in the lamina propria of the simple folds. A weak diffuse labelling was observed in the smooth muscle, which was interpreted as background labelling. The negative controls were blank. The density of CD3ε positive cells in the simple folds of groups fed diets containing SBM was increased compared with the density in FM group. IEL’s that showed labelling for CD3ε were located as individual cells at the base of the epithelium but occasionally clusters of CD3ε-labelled IEL’s were observed. CD3ε positive cells were more frequent in the lamina propria adjacent to the stratum compactum of fish fed SBM compared with fish fed FM (Fig 3D), but there were only a few CD3ε-positive cells present in the lamina propria of the simple folds (Fig 3C).

**Fig 3.**
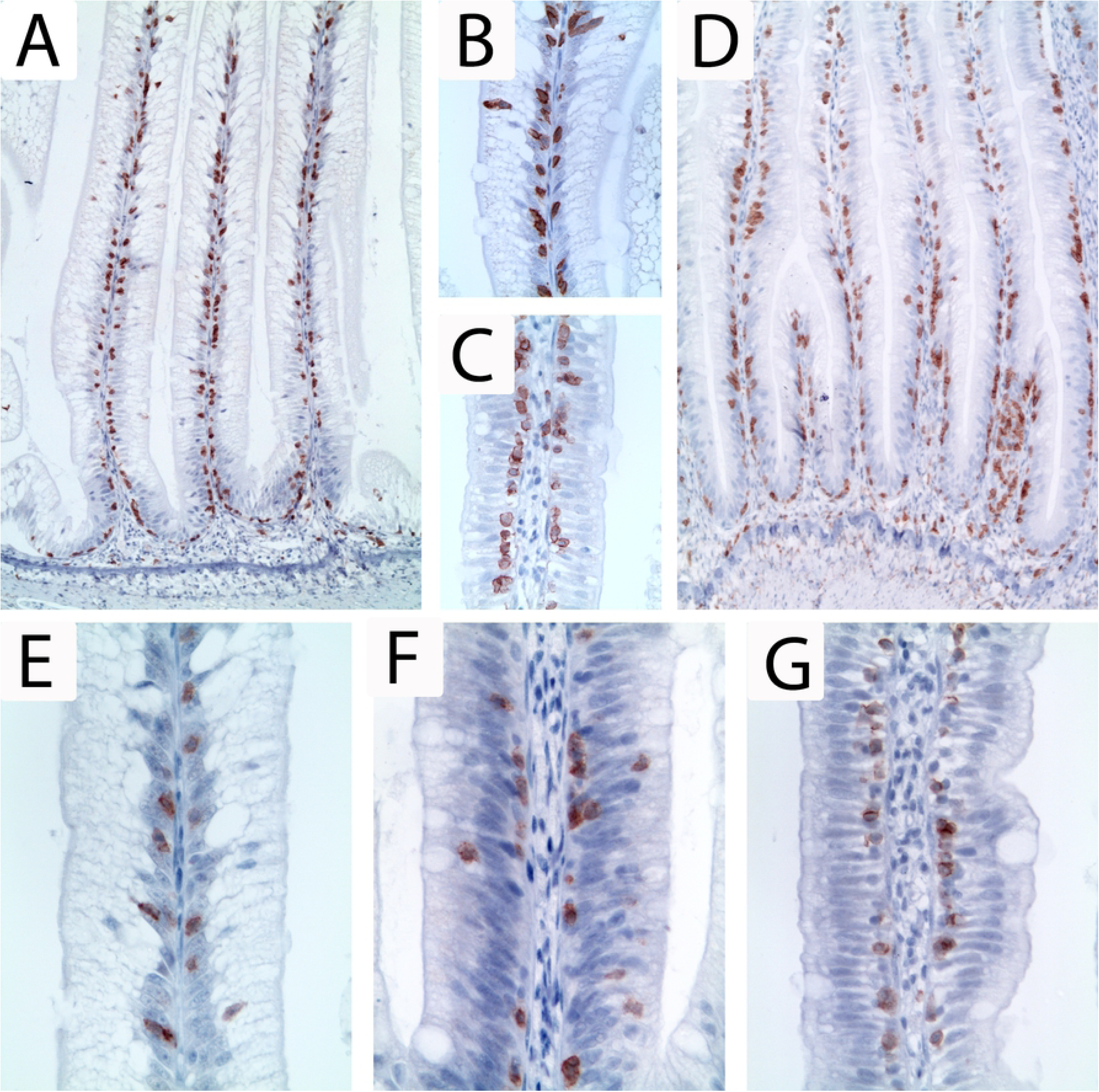
Immunohistochemical staining for CD3ε and CD8α positive cells at day 30. Immunohistochemical labelling (brown) showed an abundant presence of CD3ε positive cells at the base of the epithelium along the entire length of simple folds in all diet groups (A: FM200CU; D: SBM). At higher magnification, CD3ε positive cells were rarely present in the lamina propria of the simple folds in any of the diets (B: FM200CU; C: SBM). However, there was a higher number of CD3ε positive cells in the lamina propria adjacent to the stratum compactum in groups fed diets with SBM (D: SBM). CD8α positive cells were mainly found between the epithelial cells of all individuals of all diet groups (E: FM200CU; F: SBM200CU; G: SBM). Image A and D captured at 10x magnification, Image B, C, E, F and G captured at 40x magnification.

CD8α-labelled IEL’s showed the same distribution as the CD3ε-labelled IEL’s being located basally between the intestinal epithelial cells in all diets (Fig 3E-G). In general, the presence of CD8α-labelled IEL’s was lower than the presence of CD3ε-labelled IEL’s.

Morphometric measurements of simple folds, both length and area, revealed no significant differences between FM and FM200CU groups. The simple folds in the DI of fish fed SBM, SBM25CU and SBM200CU were significantly shorter and had a significantly smaller area than the simple folds in fish fed FM200CU and FM. There were no statistically significant differences between the simple folds of fish fed diets containing SBM either alone or combined with *C. utilis* (Fig 4A-B).

**Fig 4.**
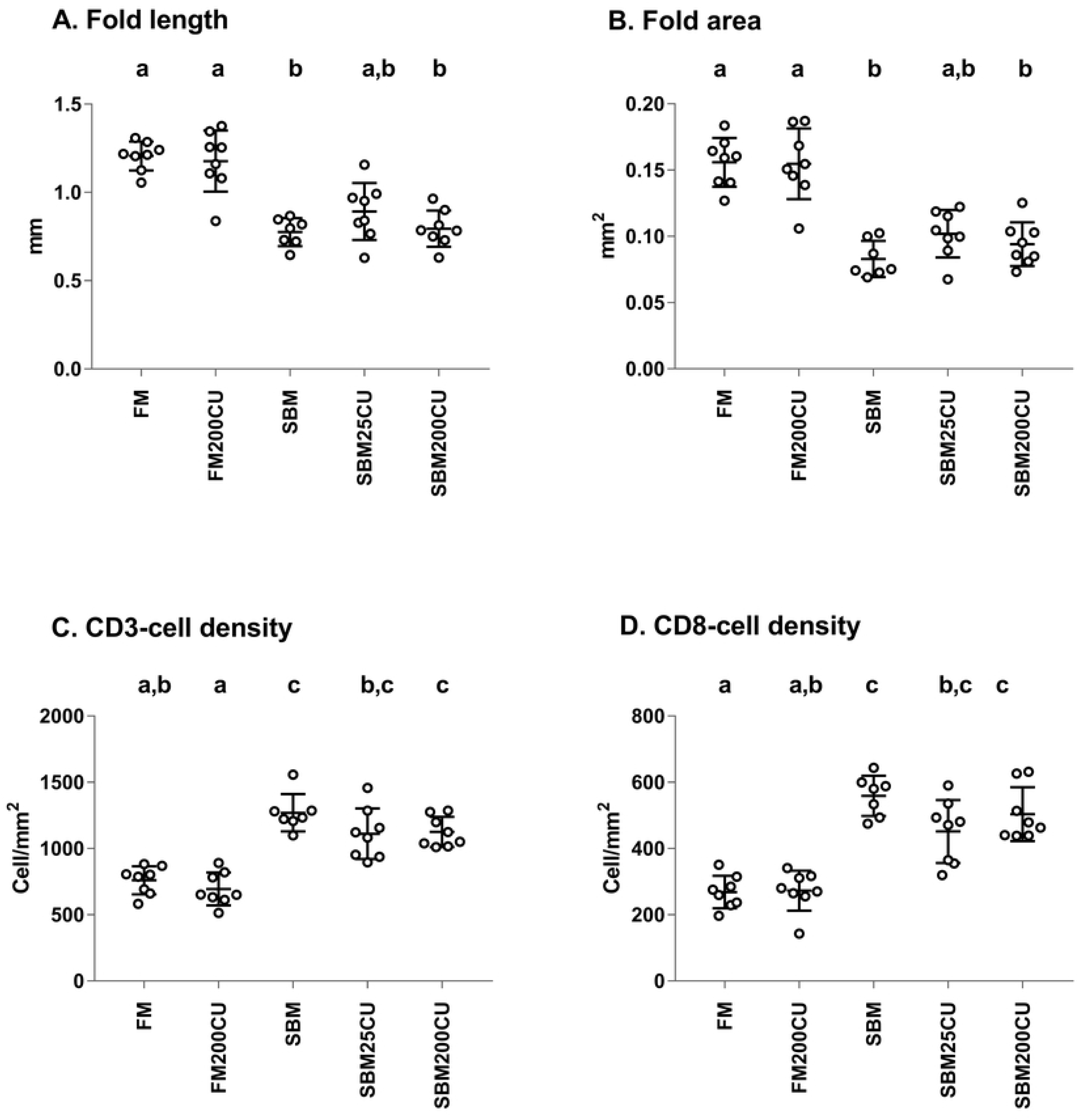
Morphometry of simple folds in distal intestine at day 30. Morphometric measurements of fold length (**A**) and fold area (**B**) of the simple folds of the distal intestine, and the density of CD3- and CD8-positive T-cells in simple folds including the lamina propria adjacent to the stratum compactum (**C** and **D**). Data are expressed as mean ± standard deviation (SD) for each individual, n = 7 for the SBM diet and n = 8 individuals per diet for the remaining groups. Groups with different letters on the upper x-axis are significantly different (p<0.05; Dunn’s test)

The density of CD3ε and CD8α-labelled cells in fish of FM and FM200CU groups was significantly lower compared with the density in fish from groups fed diets containing SBM. There was a statistically significant difference between the density of CD8α-labelled cells in fish of SBM group and the density CD8α-labelled cells in fish from the SBM25CU (p = 0.0465) (Fig 4D).

### Gene expression

Among the tested genes, only mRNA expression of Aquaporin 8 (*aqp8*) was significantly down regulated in the SBM and SBM200CU groups compared with the FM control group (Fig 5E). There was no significant difference between FM200CU and the FM control group. The transcription levels of superoxide dismutase 1 (*sod1*), glutathione S-transferase alpha 3 (*gsta3*), annexin A1 (*anxa*), and catalase (*cat*) were not different among dietary groups (p > 0.05) (Fig 5A-D).

**Fig 5.**
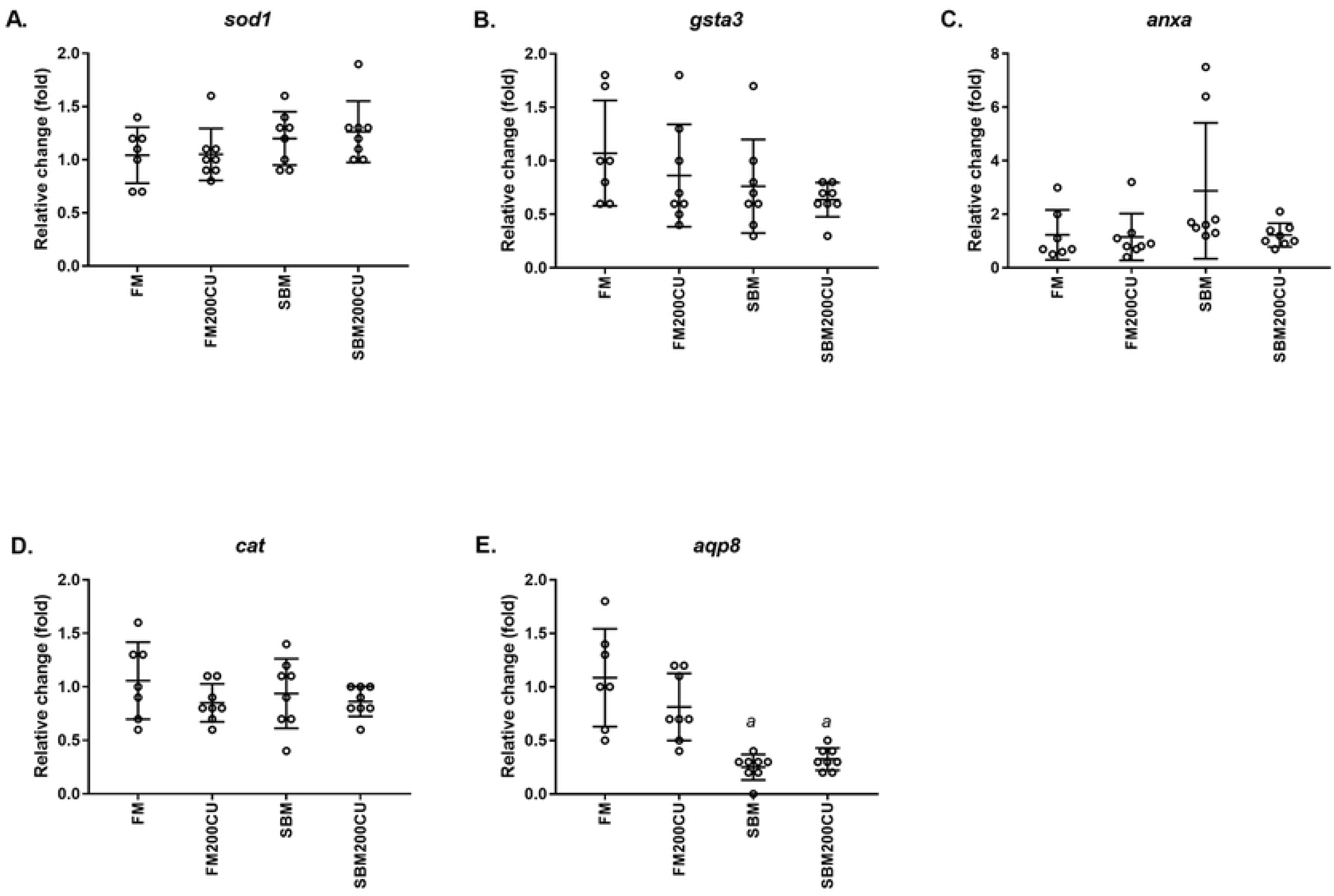
Gene expression. Quantitative PCR analyses of (**A**) superoxide dismutase 1 (*sod1*), (**B**) glutathione S-transferase alpha 3 (*gsta3*), (**C**) annexin (*anxa*), (**D**) catalase (*cat*) and (**E**) aquaporin 8 (*aqp8*) genes in the DI of Atlantic salmon fed a control fishmeal-based diet (FM), a diet containing 200 g/kg *Candida utilis* (FM200CU), and a diet containing 200 g/kg soybean meal (SBM) and one diet with 200 g/kg SBM in combination with 200 g/kg of *C. utilis* (SBM200CU) for 30 days. Data are mean –ΔΔCT ± SE (n = 7 for FM diet, n = 8 for the other groups).

### Plasma proteome

We performed proteomic analysis on plasma sampled at day 30 from four individual fish from treatment FM, SBM, SBM200CU and FM200CU. In order to simplify the reading of the data, the treatment FM is named **D1**, SBM **D2**, SBM200CU **D6** and FM200CU **D7**. In total, 367 salmon proteins were identified across the four dietary groups. A Venn diagram shows the overlap between plasma protein sets detected across the four dietary treatments (Fig 6A). There were 304 plasma proteins shared between the four groups. Moreover, each dietary treatment presented unique proteins in at least half of the replicates (Fig 6B). It is important to mention that due to the variability within the groups; just two proteins were expressed in all the four replicates in FM fed group (highlighted in Fig 6B).

**Fig 6.**
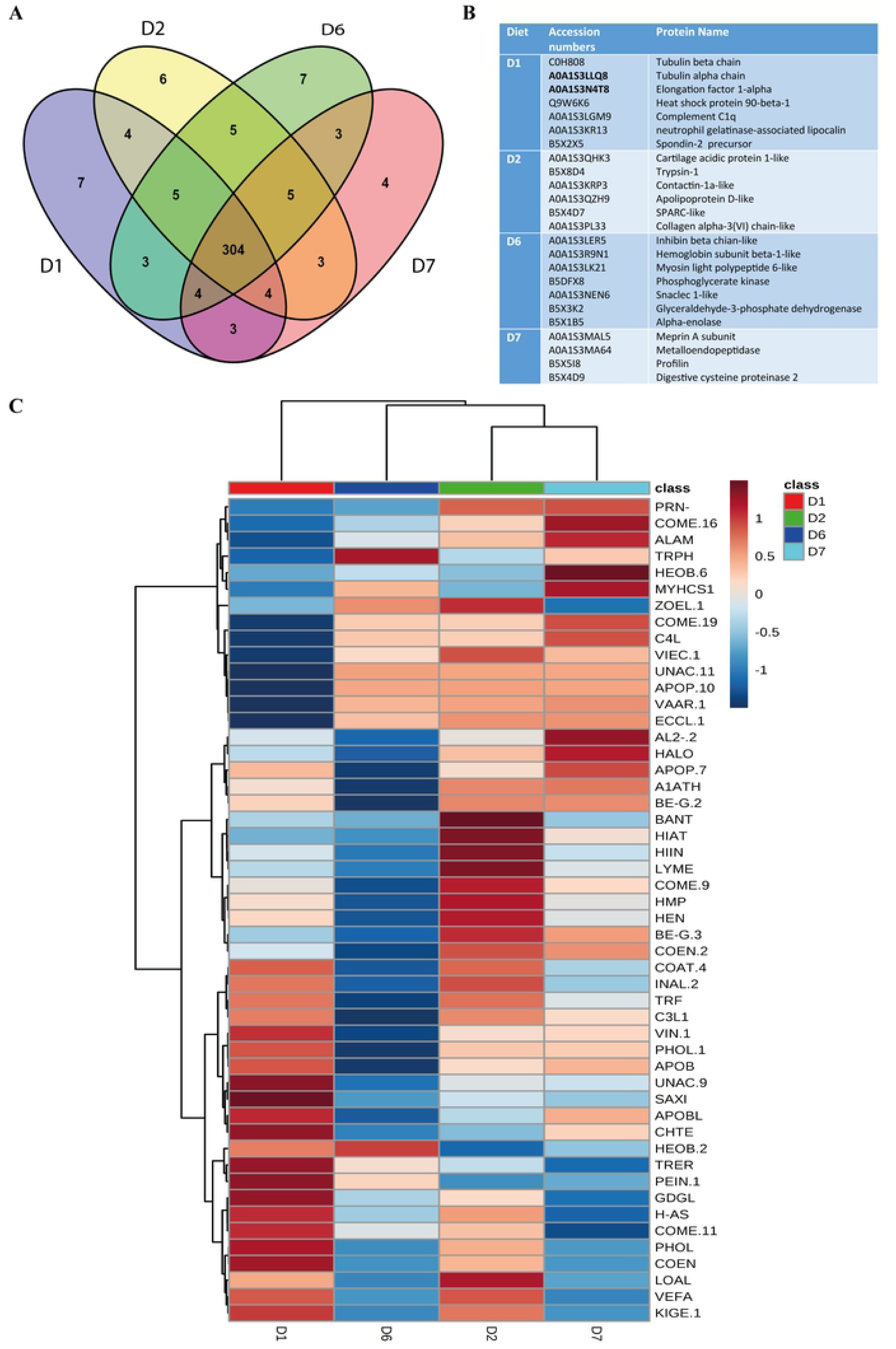
Common and unique proteins expressed in plasma of salmon fed different diets. (**A**) Venn-diagram showing the overlap between plasma protein sets detected across the four diet groups FM (D1), SBM (D2), SBM200CU (D6) and FM200CU (D7). (**B**) Unique proteins expressed in each dietary group. (**C**) Hierarchical clustering heatmap shows Spearman’s correlation between the proteins. Positive correlation values are in red, and negative correlation values are in blue.

After filtering for plasma proteins present in at least two of the four individuals per diet, 256 proteins were selected for further analyses (S3 Table). This criterion was used due to the variability among fish. To determine whether certain proteins share particular expression patterns, we produced a hierarchical clustering heatmap showing 50 significantly different proteins with positive and negative correlations based on Spearman non-parametric correlation coefficients (Spearman’s R) (Fig 6C).

The GO analysis revealed that plasma has mainly proteins involved in metabolic and cellular processes, and biological regulation (S2F Fig). GO analysis also showed that many of the plasma proteins have roles in response to stimulus and immune system processes. Further, most of the proteins belong to the signal protein domain, followed by repeat and transmembrane helix domains (S2E Fig).

PCA and PLS-DA multivariate analyses were carried out first with all selected proteins across dietary groups (Fig 7A). It is apparent that all dietary groups tend to cluster together, with the exception of the FM control group in PCA, whereas a better separation of the groups is reached by using PLS-DA (Fig 7B). The results also indicate that the resulting two principal components explain a small proportion of the total variability observed in the data, and the PLS-DA analysis offers poor prediction properties (Q2<0.5) (S2A-D Fig). This means that the resulting PCA and PLS-DA models are highly over parameterized as they used all variables. Furthermore, it also confirms that high goodness of fit (i.e. R2) can be expected in over parameterized models. The proteins (variables) that fulfilled VIP criterion (VIP>2) were determined, followed by re-running PCA and PLS-DA with the resulting 10 VIP proteins (Fig 7C).

**Fig 7.**
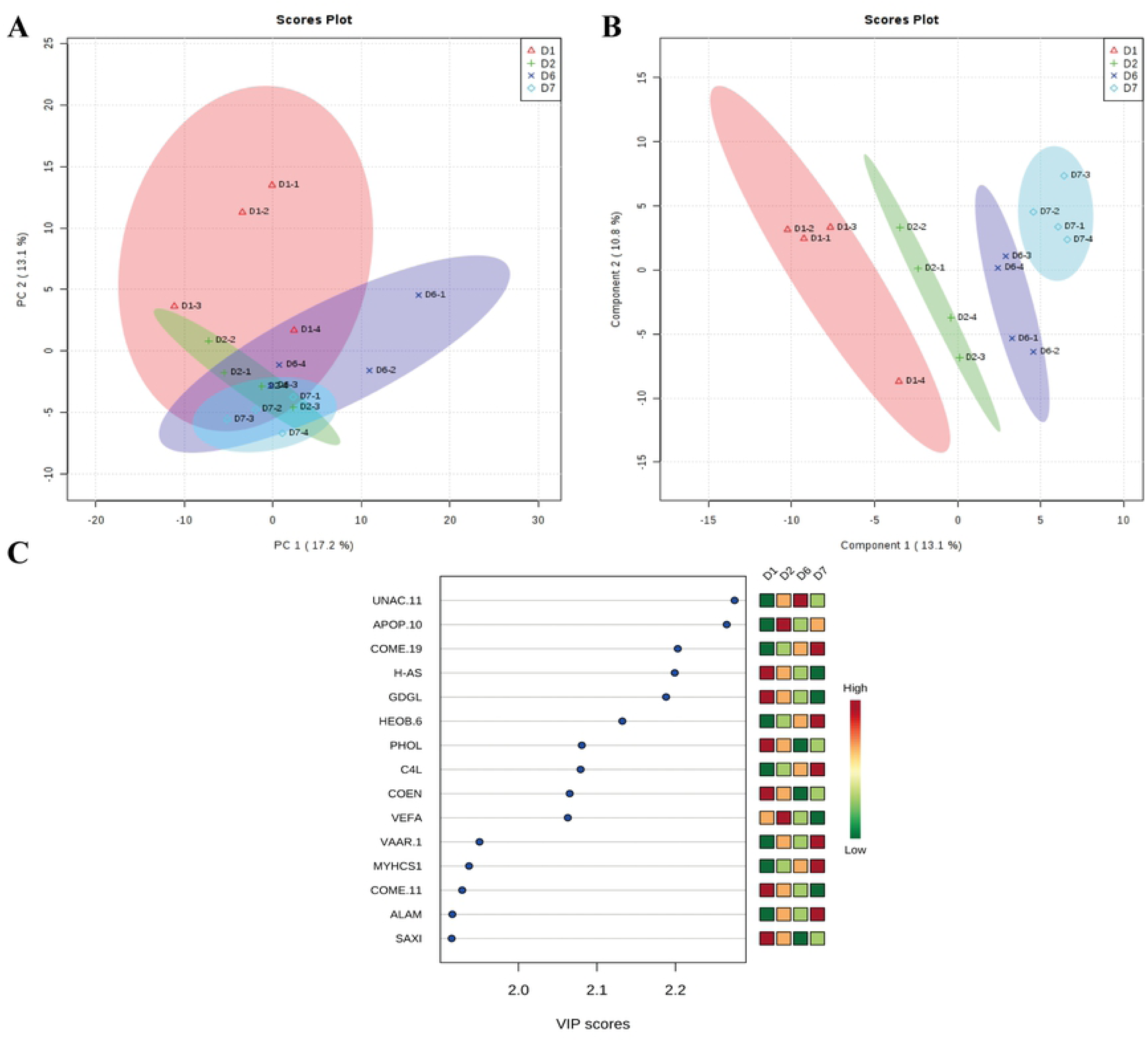
Multivariate analyses and identification of VIP proteins. **(A)** Principal components analysis (PCA) and **(B)** Partial least squares discriminant analysis (PLS-DA)’s scores plots based on selected proteins from FM (D1), SBM(D2), SBM200CU (D6) and FM200CU (D7) dietary groups. (**C**) Variable importance in the projection (VIP) scores to identify the proteins that discriminate FM (D1), SBM (D2), SBM200CU (D6) and FM200CU (D7) among the proteins selected by ANOVA plot with p value threshold 0.05.

## Discussion

Research on the effects of nutrition on fish health and disease has mainly focused on intestinal local immune responses rather than evaluating overall health impact. Therefore, the present study used an integrated approach to achieve a better understanding of the effect of feeding inactive dry *C. utilis* yeast, SBM and increasing levels of *C. utilis* yeast to Atlantic salmon in presence of SBMIE. Herein, we discuss how changes in the DI morphology and immune cell profile could reflect challenges posed by dietary treatments, while plasma protein profiling provided an association between systemic responses and outcomes of nutritional challenges.

A previous study has shown that there were no significant negative effects on feed intake, specific growth rate or feed conversion ratio when up to 30 % *C. utilis* was included in the diet for salmon [4], but DI morphology was not assessed. The present study is the first to demonstrate that FM200CU diet maintains a DI morphology similar to the FM based control diet in sea-water adapted farmed salmon. As a dietary challenge, fish were fed diets with high levels of SBM and were observed to have morphological changes in the DI that are consistent with SBMIE development in salmon [9]. Adding lower levels of *C. utilis* to SBM diets (i.e. SBM25CU and SBM50CU) resulted in a large variation within the groups, ranging from normal morphology to moderate SBMIE. The severity of SBMIE in fish fed the highest inclusion levels of *C. utilis* (i.e. SBM100CU and SBM200CU) in combination with SBM was similar to that of fish fed the SBM diet. Thus, partial prevention of SBMIE occurred only with lower inclusion levels of *C. utilis*. This finding does not agree with previous work, where feeding 200 g/kg *C. utilis* together with SBM (200 g/ kg) prevented SBMIE development in salmon [13]. This inconsistency could be due to differences in the degree of bioactivity of the *C. utilis* yeast, or in the ANF content of the SBM, and/or the experimental conditions. For example, Øverland and Skrede [11] have speculated that inconsistent effect of yeast on host immunity can be attributed to yeast strain, fermentation conditions and downstream processing when manufactured. The shift in diet at d 30 of the experiment showed that the developed enteritis observed in the DI of fish fed either SBM alone or in combination with *C. utilis* was resolved after feeding FM diet for 7 d. It is also important to point out that the degree of SBMIE were reduced from moderate to mild after 7 d feeding FM200CU diet to those fish fed SBM diet for 30 d. It can be suggested that all fish would have returned to normal state if the experiment had lasted longer.

Local DI response to SBM has been described as a T-cell mediated inflammatory response [34]. At d 30, the SBM, SBM25CU and SBM200CU groups had increased CD3ε and CD8α cell populations in the DI, which support this description. The CD3ε- and CD8α-lymphocytes were mainly confined to the basal part of the DI epithelium with only a few CD3ε-labelled cells scattered in the lamina propria adjacent to stratum compactum. However, Bakke McKellep *et al.* [34] reported that lamina propria adjacent to stratum compactum and stroma of complex folds were rich in CD3ε-labelled cells in DI presenting with SBMIE. It is relevant to indicate that there was a decreased presence of CD8α-cell population in the SBM25CU group compared with the SBM group. The CD3ε- and CD8α related observations might imply that the SBM used in these studies differed in immunostimulatory properties, and that *C. utilis* has an immunomodulating effect locally in the DI when included at a low level in the SBM diet. On the other hand, FM200CU presented a similar T-cell population profile in the DI compared with control FM group, indicating no stimulation of the T-cell population when SBMIE was not present.

The mRNA expression profile of *aqp8* gene in the DI in the diet groups (FM, FM200CU, SBM, SBM200CU) indicates an association of *aqp8* with the resulting DI morphological and immune cell responses to nutritional challenges observed in this study. Our results confirm previous findings showing suppression of *aqp8* gene expression in intestinal inflammatory processes in salmon such as SBMIE [13, 35]. The relation between *aqp8* expression and inflammation is consistent with cell-based studies showing that reduced *aqp8* expression is linked to increased oxidative cell stress damage [36] and implying that *aqp8* is a key player in the maintenance of redox cellular status. DI. In contrast, the similar mRNA *aqp8* expression levels observed in the DI from the FM200CU and FM groups can support the notion that feeding *C. utilis* to salmon promotes intestinal homeostasis. Together, *aqp8* can be considering as a good indicator to differentiate between inflamed and non-inflamed intestines in salmon.

The systemic nature of the response of salmon towards dietary treatments was pursued by the proteomic profiling of blood plasma from four diet groups at day 30 of the experiment, namely the FM, SBM, SBM200CU and FM200CU groups. By applying advanced proteomic methodologies, 10 VIP proteins and top 50 significant proteins, presented in a hierarchial clustering heatmap, visualize the changes in plasma protein levels across the dietary groups. Differences between the plasma protein profiles of fish receiving SBM diets (SBM and SBM200CU) and FM diet were expected, as systemic changes may accompany a local tissue inflammation. However, the difference in plasma protein profile between FM control group and FM200CU experimental group was noteworthy as these two dietary groups responded similarly at a local tissue level. *C. utilis* containing diets are known to have immunomodulatory effects, therefore, the proteins that are related to an immune response/function are of particular interest.

VIP protein HA-S (i.e. H-2 class I histocompatibility antigen, Q10 alpha chain) has been involved in the modulation of natural killer cells in the mouse [37], implying a role in immune tolerance [38]. Interestingly, the strongest reduction in HA-S expression was detected in the FM200CU group, which at the same time presented with DI morphology similar to the FM group. Complement factors are part of the innate immune system that enhance the ability of antibodies and phagocytic cells to clear microbes and damaged cells and promote inflammation. Previous study in Atlantic salmon have shown that intraperitoneal injection of glucans from *Saccharomyces cerevisiae* resulted in increased activities of complement in plasma 2-4 weeks after injection [39]. In our study, the FM200CU group had an increased expression of COME.19 and C4L (i.e. complement 4 like), and COME.16 (complement C4-B) as shown in the hierarchial clustering heatmap. On the other hand, the VIP protein VEFA (i.e. venom factor like) that also has a suggested role in complement activation [40] has a lower expression when compared to FM group. These findings are relevant as they highlight differences between the FM and FM200CU groups, indicating a systemic effect on the complement system by *C. utilis* without causing DI inflammation [41].

HIIN, LYME and HMP are among the significant proteins that are highly expressed in SBM, and separate the SBM group from the other groups, especially the SBM200CU group. Lysozyme (LYME) is primarily associated with innate immune defense against bacteria but is also known to activate the complement system [42]. Our results show that DI inflammation is associated with high plasma levels of LYME (i.e. Lysozyme g) in blood as seen in human with active Chron’s disease [43]. This observation also suggests that LYME expression during SBMIE in Atlantic salmon might vary depending on the tissue as mRNA down-regulation of LYME has been observed in DI with SBMIE [13]. Thus, further proteomic analysis in inflamed DI is needed to elucidate the differences between protein and gene expression analyses. Hemopexin (HMP) is regarded as an acute phase protein in mammals that possesses a high affinity for heme released from hemoglobin [44]. The sequestering of circulating heme iron is an important dual defense mechanism as bacterial growth relies on the acquisition of iron from their host [45] and free heme can cause oxidative damage [46]. Therefore, increased expression of HMP can be expected in inflammation and in infectious diseases to promote tissue healing by reducing tissue oxidative damage. This is supported by increased HMP expression during bacterial infections in Atlantic salmon [47] and rainbow trout [48]. These observations are further supported by our study showing a distinct strong HMP plasma protein expression in salmon presenting with DI inflammation induced by nutrition challenge. Fish with a high degree of DI inflammation (SBM group) also showed high HIIN (i.e. histidine-rich glycoprotein-like) levels compared with the other groups. HIIN has been shown to modulate tumor progression and anti-tumor immune responses in humans, and its high expression made it useful as a potential glycoprotein biomarker in some cancers [49]. These three proteins separate SBM from SBM200CU, which is of interest as both of these groups present with enteritis in the DI and decreased gene expression of *aqp8.*

This study utilized plasma proteomic analysis to achieve a better understanding of the effects of nutrition on fish health and disease and to identify systemic protein profiles in response to the dietary treatments. The inclusion of 200 g/kg of *C. utilis* yeast to a FM based diet (i.e. FM200CU) as a novel protein source with potential functional properties produced similar DI morphology, immune cell population and gene expression profiles, compared with a FM based control. Feeding the SBM diet induced SBMIE while feeding lower inclusion levels of *C. utilis* in combination with SBM (i.e. SBM25CU and SBM50CU) reduced the severity of SBMIE. Interestingly, higher inclusion levels of *C. utilis* yeast (i.e. SBM100CU and SBM200CU) did not show any significant protection against SBMIE, as observed in previous studies. A key finding from this study is that dietary *C. utilis* could influence the expression of plasma proteins related to immune function without altering DI morphology. Further research is needed to evaluate the impact of yeast strain and the fermentation and/or down-stream processing conditions of the yeast on functional properties in relation to gastro-intestinal health and systemic responses.

## Acknowledgments

Thanks to Aleksandra Bodura Göksu for helping with immunohistochemistry techniques and to Ricardo Tavares Benicio for helping with feed manufacture, feeding the fish and sampling.

## Supplementary material

**S1 Table. Chemical composition (g kg-1) of dry *Candida utilis* biomass.**

**S2 Table. Primers used in qPCR analysis.**

**S3 Table. List of 256 proteins identified**

**S1 Fig. Morphometric measurement.** Red line indicates measurement of fold height from the tip of the simple fold to the stratum compactum. The yellow line indicates the fold area including the simple fold and the lamina propria adjacent to the stratum compactum.

**S2 Fig. Multivariate analyses and Gene ontology analyses of plasma proteins.** Plots **(A)** and **(B)** shown R2 (goodness of fit) and Q2 (goodness of prediction) for the PLS-DA models before and after discrimination of VIP variables. The red star indicates the best classifier. The explained variances are shown in brackets. The explained variances are shown in brackets. Score plots **(C)** and **(D)** shown PCA and PLS-DA based on VIP variables. (**E**) Protein domain analysis reveals that plasma is enriched in proteins with signal domain, repeat and coiled domain. **(G)** Gene ontology analysis reveals that plasma is enriched in proteins with functions in metabolic and cellular processes and biological regulation. The numbers of proteins in each Gene Ontology class are shown.

